# Adiponectin signaling modulates fat taste responsiveness in mice

**DOI:** 10.1101/2024.09.05.611494

**Authors:** Fangjun Lin, Emeline Masterson, Timothy A. Gilbertson

**Affiliations:** Burnett School of Biomedical Sciences, College of Medicine, University of Central Florida, Orlando, FL 32827, USA; Department of Internal Medicine, College of Medicine, University of Central Florida, Orlando, FL 32827, USA

## Abstract

We previously reported that the adiponectin receptor agonist AdipoRon selectively enhances cellular responses to fatty acids in a human taste cell line. The enhancement role of AdipoRon on fatty acid-induced cell responses is mediated by the activation of AMPK and translocation of CD36 on human taste cells. It has also been shown that adiponectin selectively increases taste behavioral responses to intralipid in mice. However, the molecular mechanism underlying the physiological effects of adiponectin on fat taste in mice remains unclear. Here we define AdipoR1 as the mediator responsible for the enhancement role of adiponectin/AdipoRon on fatty acid-induced responses in mouse taste bud cells. Calcium imaging data demonstrate that AdipoRon enhances linoleic acid-induced calcium responses in a dose-dependent fashion in mouse taste cells isolated from circumvallate and fungiform papillae. Similar to the human taste cells, the enhancement role of AdipoRon on fatty acid-induced responses was impaired by the co-administration of an AMPK inhibitor (Compound C) or a CD36 inhibitor (SSO). Utilizing Adipor1-deficient animals we determined the enhancement role of AdipoRon/adiponectin is dependent on AdipoR1 since AdipoRon/adiponectin failed to increase fatty acid-induced calcium responses in taste bud cells isolated from these mice. Brief-access taste tests were performed to determine whether AdipoRon’s enhancement role was correlated with any differences in taste behavioral responses to fat. Although AdipoRon enhances the cellular responses of taste bud cells to fatty acids, it does not appear to alter fat taste behavior in mice. However, fat naïve *Adipor1*^−/−^ animals were indifferent to increasing concentrations of intralipid, suggesting that adiponectin signaling may have profound effects on the ability of mice to detect fatty acids in the absence of previous exposure to fatty acids and fat-containing diets.

## Introduction

Adiponectin, a hormone derived from adipose tissue, acts as a messenger between fat tissue and other organs and plays essential roles in regulating energy homeostasis, obesity, diabetes, cardiac function, and inflammation. For example, adiponectin inhibits gluconeogenesis in the liver and enhances fatty acid oxidation in skeletal muscle, contributing to beneficial metabolic effects in energy homeostasis (Yamauchi et al., 2002). Adiponectin can improve endothelial dysfunction caused by elevated free fatty acids (Li et al., 2007; Wang et al., 2012). Adiponectin protects against obesity-related glomerulopathy and diabetic nephropathy (Xu et al., 2021; Yi & OuYang, 2019). Moreover, adiponectin has also been found to enhance 5’ adenosine monophosphate-activated protein kinase (AMPK) activity in the hypothalamus, stimulating food intake (Kubota et al., 2007). Adiponectin could also be produced by salivary gland epithelial cells (Katsiougiannis et al., 2006) and the presence of salivary adiponectin has been identified in several studies (Akuailou et al., 2013; Nigro et al., 2015; Toda et al., 2007). In addition, high expression levels of adiponectin receptors were observed in mouse taste bud cells (Crosson et al., 2019) and immortalized human fungiform (HuFF) taste cells (Lin et al., 2023). Thus, these studies suggest a potential role for adiponectin signaling in taste perception.

There is growing evidence that taste function can be modulated by hormones that act on their receptors in the peripheral gustatory system. Of the many hormones that influence taste perception, several appear to modulate the cellular, neural, or behavioral responses to fatty acids, such as leptin, peptide YY (PYY), glucagon-like peptide-1 (GLP-1), ghrelin, cannabinoids, and adiponectin. For example, leptin suppresses cellular and neural responses to fatty acids (Kitagawa et al., 2007; Ullah et al., 2021). The behavioral responses to lipid emulsions were attenuated in *PYY* knockout mice, and it was rescued by the targeted augmentation of salivary PYY via viral vector therapy (La Sala et al., 2013). Administration of the GLP-1 receptor agonist, exendin-4, decreased taste responses to intralipid (Treesukosol & Moran, 2022), and *Glp1r* knockout mice lost the ability to detect low concentrations of lipids (Martin et al., 2012). Reduced taste sensitivity to intralipid was observed in *ghrelin* and ghrelin O-acyltransferase (*GOAT*) knockout mice (Cai et al., 2013), and ghrelin receptor (growth hormone secretagogue receptor, *Ghsr*) knockout females but not males showed a reduction in taste responsiveness to linoleic acid (Calder et al., 2021). Cannabinoid 1 receptor (*Cb1r*)-deficient mice displayed a low preference for fatty solutions compared to wild-type mice (Brissard et al., 2018). Moreover, salivary gland-specific adiponectin rescue significantly increased the behavioral taste responses to intralipid in adiponectin knockout mice (Crosson et al., 2019). We recently reported that adiponectin selectively enhances fatty acid-induced calcium responses in immortalized human fungiform taste cells by increasing the surface expression of CD36 (Lin et al., 2023). However, the physiological connection between adiponectin signaling and fat taste sensitivity in the native taste system, if any, is largely unexplored.

Adiponectin acts mainly on three receptors, including AdipoR1, AdipoR2, and T-cadherin. These receptors are widely distributed throughout the body, with AdipoR1 most abundantly expressed in skeletal muscle and AdipoR2 predominantly expressed in the liver (Yamauchi et al., 2003). Immunohistochemical staining showed that both AdipoR1 and T-cadherin immunolocalize to mouse circumvallate taste buds, while AdipoR2 immunolocalize to the surrounding tissues, but not in taste buds (Crosson et al., 2019). AdipoR1 and AdipoR2 mediate different pathways of adiponectin downstream signaling and show opposing effects on energy metabolism in mice (Bjursell et al., 2007). AdipoR1 appears to be more tightly associated with the activation of the AMPK pathway, whereas AdipoR2 is more closely involved in the activation of the peroxisome proliferator-activated receptor (PPAR)α pathway (Yamauchi et al., 2007). Additionally, our previous study showed that the EC_50_ of AdipoRon’s enhancement of linoleic acid (LA)-induced calcium response is close to the reported K_d_ of AdipoRon binding to AdipoR1 and that the effect of AdipoRon on LA-induced calcium responses in HuFF cells requires the activation of AMPK (Lin et al., 2023). We hypothesized that AdipoR1 may be responsible for the role of adiponectin on fat taste responsiveness. To explore this hypothesis, we first aimed to validate if AdipoRon/adiponectin could enhance the fatty acid-stimulated calcium responses in mouse taste bud cells and whether AMPK and CD36 are required for this enhancement role. Next, we examined the contribution of AdipoR1 using taste bud cells isolated from *Adipor1* gene knockout mice. Our results indicate that AdipoRon/adiponectin increased taste responses to fatty acids in taste cells through the activation of AdipoR1 and AMPK. Following the functional studies in these cell-based assays, we attempted to investigate the role of AdipoRon in regulating fat taste behavior in mice. The data showed that the concentration of AdipoRon that enhanced the cellular responses of taste bud cells to fatty acids did not appear to alter the animals’ behavior toward the fatty acids. Lick responsiveness in brief-access taste testing was unchanged by directly adding the AdipoRon (10 μM) into the testing solutions. Moreover, brief-access taste testing in *Adipor1*-deficient mice showed that naïve *Adipor1*-deficient male mice failed to taste the intralipid solutions, while naïve *Adipor1*-deficient female animals could detect the difference between intralipid and water but were indifferent to increasing concentrations of intralipid. These findings suggest that adiponectin signaling has a profound, possibly sex-dependent, function in regulating fat taste.

## Materials and Methods

### Animals

The mouse strain (B6.129P2-*Adipor1*^tm1Dgen^/Mmnc, RRID: MMRRC_011599-UNC) was donated to the NIH-sponsored Mutant Mouse Resource and Research Center (MMRRC) by Deltagen. Heterozygous *Adipor1*^+/−^ mice were ordered from the MMRRC facility at the University of North Carolina. Adult C57BL/6J mice were obtained from Jackson laboratories or the Lake Nona animal facility at the University of Central Florida. Mice were housed under standard laboratory conditions (12 h:12 h day/night cycle) with water and standard chow available *ad libitum* unless otherwise specified. All animal procedures were approved by the Institutional Animal Care and Use Committee of the University of Central Florida.

### Genotype and Phenotype of the Adipor1-deficient Mice

At about two weeks of age, newborn mice were ear-punched. Genomic DNA was extracted from ear punch samples with DirectPCR Lysis Reagent (Ear) and Proteinase K (Viagen Biotech, Los Angeles, CA, USA). Polymerase chain reaction (PCR) was performed with primers (sequences 5’ to 3’) as follows: NIH62-GS1, TCCACTGTGTCAGCTTCTCTGTTAC, NIH62-GS2, AGGCAGGGTAAGCTGATTAGCTATG, and NIH62-neo, GGGTGGGATTAGATAAATGCCTGCTCT (MMRRC Center Protocol 11599, www.med.unc.edu/mmrrc). The PCR cocktail contained 1 μl crude ear lysates, 1 μl 20 μM primers mixture, 6 μl MyTaq™ Red Mix (Meridian Bioscience, Memphis, TN, USA), and 5 μl water. The PCR was then carried out according to the genotyping protocol 11599 with initial heating to 94°C for 5 min, then 36 cycles of 94°C for 45 sec, 60°C for 45 sec, and 72°C for 90 sec, with a final extension at 72°C for 7 min. After electrophoretic separation on 2% agarose gels, the expected sizes of bands (254 bp for wild type and 433 bp for mutant) were visualized with ChemiDoc MP Imaging System (Bio-Rad, Hercules, CA, USA).

Body weights were recorded at about six weeks of age, and total body composition (fat mass, lean mass, and free body fluid) was determined by using the Bruker time domain-nuclear magnetic resonance (TD-NMR, minispec LF50) live mice body composition analyzer (Bruker, Billerica, MA, USA). The homozygous *Adipor1* knockout (−/−) mice appeared normal at birth and subsequently developed and bred normally. Both the males and females of the *Adipor1*^−/−^ mice were fertile. At about six weeks of age, the mean body weight of eight *Adipor1*^+/+^ male mice was 23.17 g, and that of eight *Adipor1*^+/+^ female mice was 18.42 g, while that of seven *Adipor1*^−/−^ male mice was 22.07 g and that of nine *Adipor1*^−/−^ female mice was 17.53 g (Figure S1A). The differences of the total body composition between *Adipor1*^+/+^ and *Adipor1*^−/−^ mice in males and females were shown in Figure S1B-D.

### Chemicals and Solutions

Standard Tyrode’s solution contained (in mM) 140 NaCl, 5 KCl, 1 CaCl_2_, 1 MgCl_2_, 10 HEPES, 10 glucose, and 10 Na pyruvate; adjusting the pH to 7.40 with NaOH; 300–320 mOsm. Calcium-magnesium free Tyrode’s solution contained (in mM): 140 NaCl, 5 KCl, 2 BAPTA, 10 HEPES, 10 glucose, and 10 Na pyruvate; adjusting the pH to 7.40 with NaOH; 300–320 mOsm. Linoleic acid (LA) was purchased from Sigma (St. Louis, MO, USA), prepared as stock solutions (25 mg/ml) in 100% ethanol, and then stored under nitrogen at −20 °C. LA working solutions were made from stock solutions immediately before use and only used on that day. Intralipid 20% IV fat emulsion was purchased from Patterson Veterinary (Loveland, CO, USA). Isolation enzyme cocktail components: collagenase A (0.5 mg/ml), dispase II (2 mg/ml), and trypsin inhibitor (1 mg/ml) were ordered from Sigma. Stock solutions of AdipoRon (MedChem Express, Monmouth Junction, NJ, USA), sulfosuccinimidyl oleate (SSO; Cayman Chemical, Ann Arbor, MI, USA), and dorsomorphin (Compound C; ApexBio, Houston, TX, USA) were made in DMSO and diluted the day of the experiment to a designated concentration. The R&D systems™ recombinant mouse adiponectin protein was ordered from Fisher Scientific.

### Taste Cell Isolation

Individual taste bud cells were isolated from mouse circumvallate and fungiform papillae following procedures used in previous reports. Briefly, adult males and females from both *Adipor1*^−/−^ mice and *Adipor1*^+/+^ controls were sacrificed by exposure to CO_2_ followed by cervical dislocation. Tongues were removed and immediately placed in Tyrode’s solution. The anterior portion of the tongue containing the fungiform papillae and the area surrounding the circumvallate papillae were then injected with an enzyme cocktail in Tyrode’s solution. The injected tongue was incubated in Tyrode’s solution and bubbled with O_2_ for approximately 40 minutes. The lingual epithelium was separated from the underlying muscle layer and pinned flat in Tyrode’s solution in a Sylgard-coated petri dish. Next, the epithelium was incubated in calcium-magnesium free Tyrode’s solution for 5–7 minutes (males) and 3–5 minutes (females), washed with standard Tyrode’s solution, and subsequently in the same enzyme cocktail described above for another 3–5 minutes (males) and 1–3 minutes (females). Taste bud cells were isolated by gentle suction using a fire-polished glass pipette under a dissection microscope and placed on coverslips coated with Corning Cell-Tak Cell and Tissue Adhesive (Corning, PA, USA) for calcium imaging.

### Calcium Imaging

Intracellular calcium imaging was carried out on isolated taste bud cells loaded with 4 µM of the ratiometric calcium indicator Fura-2-acetoxymethyl ester (Fura-2 AM, Invitrogen) in Tyrode’s with 0.05% pluronic acid F-127 (Invitrogen) for about 1 hour at room temperature in the dark. The coverslips with isolated taste bud cells ready for imaging were placed onto a perfusion chamber (RC-25F, Warner Instruments, Holliston, MA, USA). Taste stimuli solutions (30 μM LA with or without other compounds) were applied by a bath perfusion system at a flow rate of 4 mL/minute for 3 minutes, followed by 1 minute of 0.1% fatty acid-free BSA solution, and then regular Tyrode’s for about 2 minutes to remove the LA until the calcium signal returned to near baseline level. Taste bud cells were illuminated with Lambda DG-5 illumination system (Sutter Instruments, Novato, CA, USA), and imaging was performed using an acA720 camera (Basler, Ahrensburg, Germany) through a 40× oil immersion objective lens of an Olympus CKX53 inverted microscope. Taste bud cells loaded with Fura-2 AM were excited at 340 nm and 380 nm of light, and emission was recorded at 510 nm. Images were captured at a rate of 20 per minute, and the ratio of fluorescence (340 nm/380 nm) was used to measure the changes in intracellular calcium levels on taste bud cells by InCyt Im2™ imaging software (Version 6.00, Cincinnati, OH, USA). Experiment 1: To observe the effect of AdipoRon on fatty acid-induced calcium responses in isolated taste bud cells from wild-type mice, LA (30 μM) and mixtures of LA (30 μM) with a series of AdipoRon concentrations (0.1, 1, 5, and 10 μM) were used. Experiment 2: LA (30 μM), a mixture of LA (30 μM) with AdipoRon (5 μM), and a mixture of LA (30 μM), AdipoRon (5 μM) with Compound C (an AMPK inhibitor, 10 μM) in random order were perfused over isolated wild-type mouse taste bud cells to link the activation of AMPK to the enhancement role of AdipoRon on fatty acid-induced calcium responses. Experiment 3: To investigate the involvement of CD36 in the effect of AdipoRon/adiponectin on fatty acid-induced responses, LA (30 μM) and mixtures of LA (30 μM) with AdipoRon (5 μM) or adiponectin (10 ng/ml) were perfused in random order over isolated wild-type mouse taste bud cells pretreated with SSO (an irreversible inhibitor of CD36, 100 µM, 30 min). Experiment 4: To determine whether the effect of AdipoRon/adiponectin on fatty acid-induced calcium responses is mediated by AdipoR1, LA (30 μM), and mixtures of LA (30 μM) with AdipoRon (5 μM) or adiponectin (10 ng/ml) were perfused in random order over taste bud cells isolated from *Adipor1*^−/−^ mice and *Adipor1*^+/+^ controls.

### Brief-access taste testing

Four groups of animals (*Adipor1*^+/+^ males, *Adipor1*^+/+^ females, *Adipor1*^−/−^ males, and *Adipor1*^−/−^ females) were used for the brief-access testing. Mice were housed individually in standard cages and habituated to their environment for several days before testing began. Purified water was used to prepare test solutions. The brief-access testing was administered within an MS-160 Davis Rig gustatory behavioral apparatus (Med Associates, St. Albans, VT, USA). Water was restricted for about 21 hours before each training and testing day (30-minute test in Davis Rig, 1 hour rest period, then free access to water for 1.5 hours). Animals were trained by two steps: (1) one presentation of purified water for 15 min and a 15-minute time limit to the first lick on the first day; (2) on days 2-3, training and testing on a series of sucrose solutions (25, 50, 100, and 200 mM) consisted thirty 5 second presentations with 7.5-second intertrial intervals. Following the training phase, mice were subjected to five days of testing with intralipid solutions (0.5, 1, 5, 10, and 20%). The test sessions were 30 minutes, during which animals could initiate as many trials as possible. AdipoRon (10 μM) was added on testing days 3 and 4 to investigate its role in taste behavioral responses to intralipid.

### Statistical Analysis

Calcium imaging data analyses were based on the amplitude of the intracellular calcium concentration and analyzed in Origin 9.6 (Version 9.6.0.172, OriginLab, Northampton, MA, USA). Due to the water deprivation, animals licked water or tastants profusely at the beginning of the brief-access taste testing. Once they had quenched their initial thirst, the number of water licks dropped dramatically compared to the preferred stimuli. To better represent the fundamental differences in taste behavioral responses between water and tastant, trials were included when the number of water licks began to drop dramatically. The tastant/water lick ratio was calculated by dividing the mean number of licks per trial for each concentration by the mean number of water licks per trial for that animal. Statistical analysis was performed using GraphPad Prism 10 (Version 10.2.1 (395), GraphPad Software, Boston, MA, USA). Unpaired Student’s t-test, one-way ANOVA, and two-way ANOVA were used as appropriate in each experiment. The level of significance was set at α = 0.05 for all experiments. All data are presented as mean ± SEM. For most bar graphs, individual values were not plotted due to the large number of points and conditions that made the graphs virtually uninterpretable visually.

## Results

### AdipoRon enhances cellular responses to fatty acids in mouse taste cells

We recently reported that adiponectin receptor agonist, AdipoRon, enhances fatty acid-induced calcium responses in a human fungiform (HuFF) cell line (T0029; Applied Biological Materials, Richmond BC). To validate if there is a similar enhancement role of AdipoRon on calcium responses to fatty acids in isolated native mouse taste bud cells, we examined the intracellular calcium levels in isolated circumvallate and fungiform taste bud cells stimulated by 30 μM LA and 30 μM LA with different concentrations of AdipoRon (0.1, 1, 5, and 10 μM). Similar to our results in HuFF cells, AdipoRon enhanced the 30 µM LA-evoked calcium responses in a dose-dependent manner, with little to no effect seen below 1 μM and an obvious enhancement above 5 μM in isolated fungiform and circumvallate taste bud cells from both male and female mice (Figure 1A-D, n = 30−197 cells per point from at least 3 animals for each group). Unpaired, two-tailed Student’s t-test (with Welch’s correction if the standard deviations of the populations are not equal) was used to determine statistical significance (*, P < 0.05; **, P < 0.01; ***, P < 0.001 compared to the LA alone control group).

**Figure 1.**
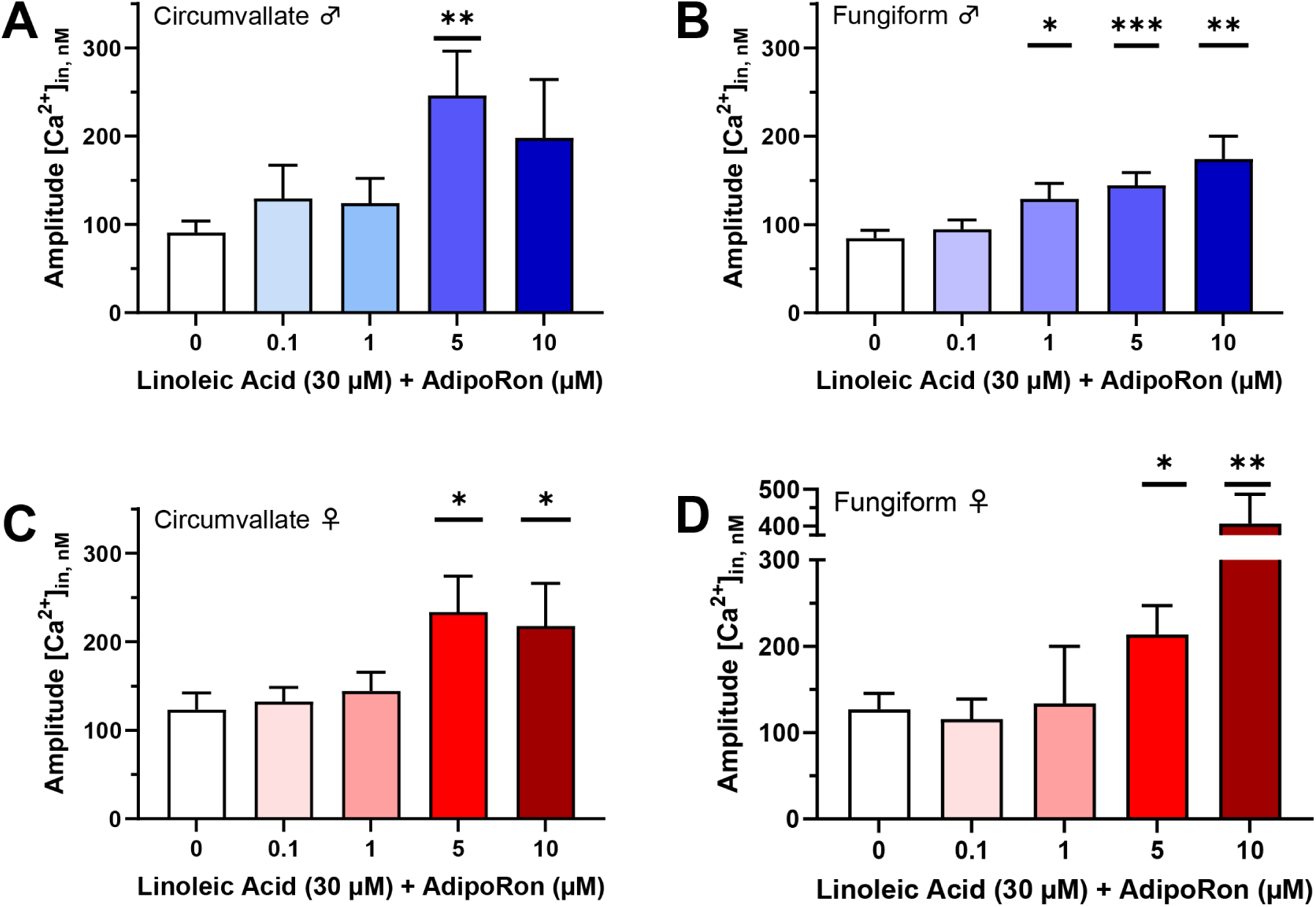
Mean amplitude of calcium responses to 30 µM linoleic acid (LA) with or without different concentrations of AdipoRon in mouse taste bud cells. (A) LA-induced calcium response was significantly increased by 5 µM AdipoRon in circumvallate taste bud cells from male mice (t_(78.53)_ = 2.995, P=0.0037); (B) LA-induced calcium response was significantly enhanced by 1 (t_(206.9)_ = 2.253, P=0.0253), 5 (t_(331.1)_ = 3.49, P=0.0005), and 10 µM (t_(77.13)_ = 3.289, P=0.0015) AdipoRon in fungiform taste bud cells from male mice; (C) LA-induced calcium response was significantly increased by 5 (t_(109.2)_ = 2.468, P=0.0151) and 10 µM (t_(126)_ = 2.114, P=0.0365) AdipoRon in circumvallate taste bud cells from female mice; (D) LA-induced calcium response was significantly enhanced by 5 (t_(152.7)_ = 2.27, P=0.0246), and 10 µM (t_(50.06)_ = 3.409, P=0.0013) AdipoRon in fungiform taste bud cells from female mice. Data are presented as mean ± SEM. Unpaired, two-tailed Student’s t-test (with Welch’s correction if the standard deviations of the populations are not equal) was used to determine statistical significance for the LA response compared with the treatments of AdipoRon. * p < 0.05, ** p < 0.01, *** p < 0.001.

### The effect of AdipoRon on LA-induced responses is mediated by AMPK activation in mouse taste cells

AdipoRon increases the phosphorylation state of AMPKα (Thr^172^) in HuFF cells, and AdipoRon’s ability to enhance LA-induced response is associated with the activation of AMPK (Lin et al., 2023). Therefore, we used a pharmacological approach to test whether activation of AMPK mediates the enhancement role of AdipoRon on fatty acid-induced calcium responses in isolated mouse taste bud cells in a similar fashion. The enhancement of taste cell responses to LA (30 μM) caused by the application of AdipoRon (5 μM) was effectively eliminated by the addition of AMPK inhibitor Compound C (10 μM). This acute response was found in both circumvallate and fungiform taste papillae cells isolated from male and female mice (Figure 2A, male circumvallate, F*_(2, 51.63)_ = 10.97, P = 0.0001, W_(2, 58.43)_ = 8.772, P = 0.0005; 2B, male fungiform, F_(2, 105)_ = 3.562, P = 0.0319; 2C, female circumvallate, F*_(2, 48.23)_ = 11.35, P < 0.0001; W_(2, 60.16)_ = 6.545, P = 0.0027; 2D, female fungiform, F*_(2, 137.6)_ = 8.508, P = 0.0003; W_(2, 103.3)_ = 7.086, P = 0.0013). A one-way ordinary ANOVA (assuming equal standard deviations) with Tukey’s test for multiple comparisons was used to determine statistical significance. However, the P values for the tests of equal variances based on the data from male circumvallate, female circumvallate, and female fungiform are minimal, the Brown-Forsythe and Welch ANOVA tests (do not assume that all standard deviations are equal) were used to determine statistical significance in these experiments instead.

**Figure 2.**
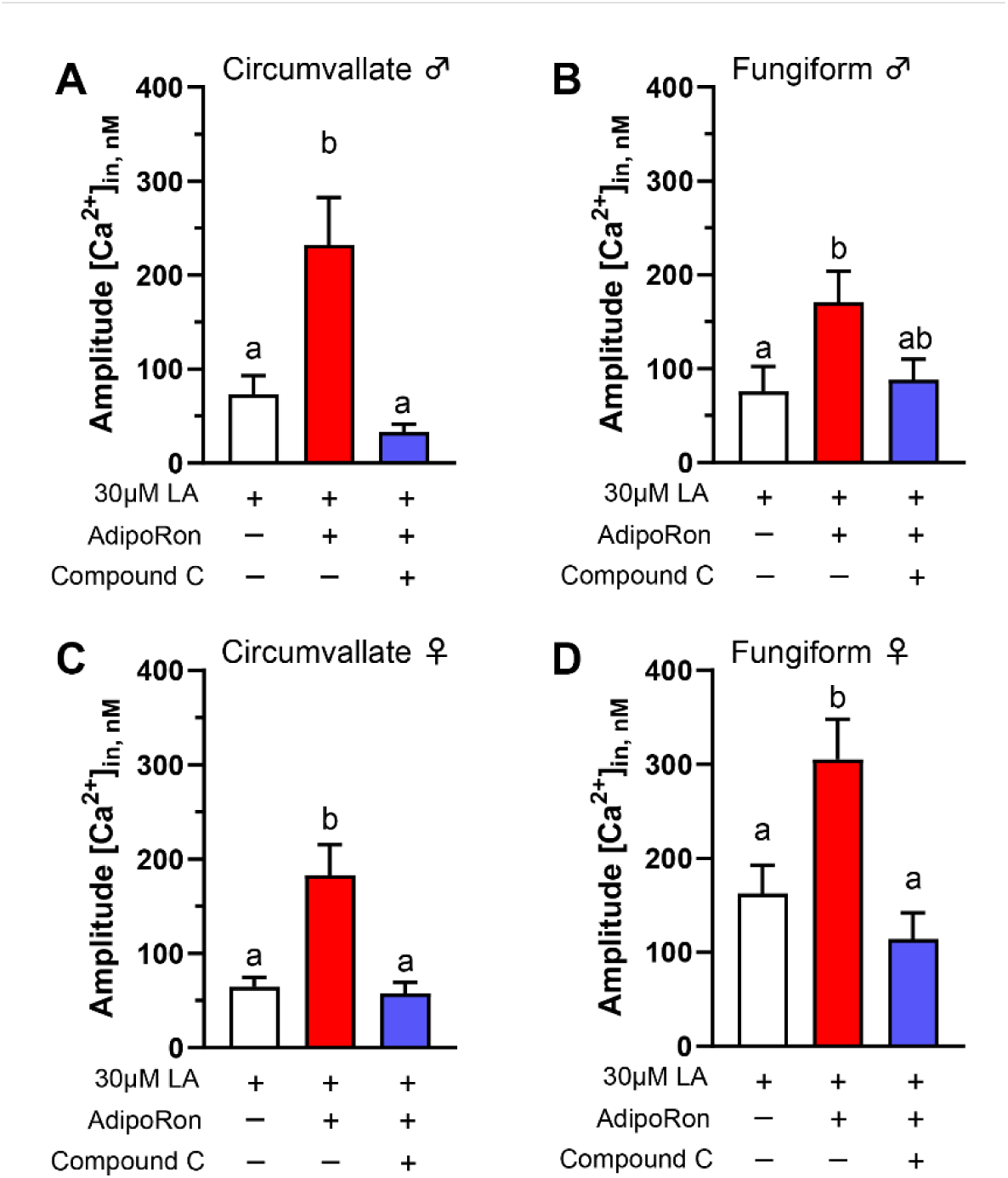
AMPK inhibitor (Compound C) eliminates AdipoRon’s ability to enhance LA-induced calcium responses in mouse taste bud cells. The enhancement of taste cell responses to 30 μM LA caused by application of 5 μM AdipoRon was effectively eliminated by the addition of 10 μM Compound C in mouse taste bud cells from (A) male circumvallate (F*_(2, 51.63)_ = 10.97, P=0.0001; W_(2, 58.43)_ = 8.772, P=0.0005), (B) male fungiform (F_(2, 105)_ = 3.562, P=0.0319), (C) female circumvallate (F*_(2, 48.23)_ = 11.35, P<0.0001; W_(2, 60.16)_ = 6.545, P=0.0027), and (D) female fungiform (F*_(2, 137.6)_ = 8.508, P=0.0003; W_(2, 103.3)_ = 7.086, P=0.0013) taste cells. Data are presented as mean ± SEM. A one-way ordinary ANOVA with Tukey’s test for multiple comparisons was used to determine statistical significance with equal standard deviations. Brown-Forsythe and Welch ANOVA tests with Dunnett’s T3 multiple comparisons test were introduced when standard deviations significantly differed. Letters above bars indicate statistical grouping.

### Adiponectin/AdipoRon acts on CD36 to increase LA-induced responses in mouse taste cells

In HuFF cells, we found that AdipoRon’s effect on the enhancement of fatty acid-induced calcium responses is mediated by CD36 signaling but not GPR120 signaling. Here, we performed a pharmacological approach to investigate the involvement of CD36 in the effect of AdipoRon/adiponectin on fatty acid-induced responses. Mouse taste bud cells were pre-incubated with a CD36 irreversible inhibitor SSO (100 µM) for 30 min. Following incubation, cells were perfused LA (30 μM) and mixtures of LA (30 μM) with AdipoRon (5 μM) or adiponectin (10 ng/ml) in random order. As shown in Figure 3, no significant differences in calcium responses were observed in SSO-pretreated taste bud cells stimulated by LA with or without AdipoRon or adiponectin (Figure 3A-D; P > 0.05 for all tests, n = 44−71 cells from at least 3 animals for each group). Unpaired, two-tailed Student’s t-test (with Welch’s correction if the populations’ standard deviations are not equal) was used to determine statistical significance.

**Figure 3.**
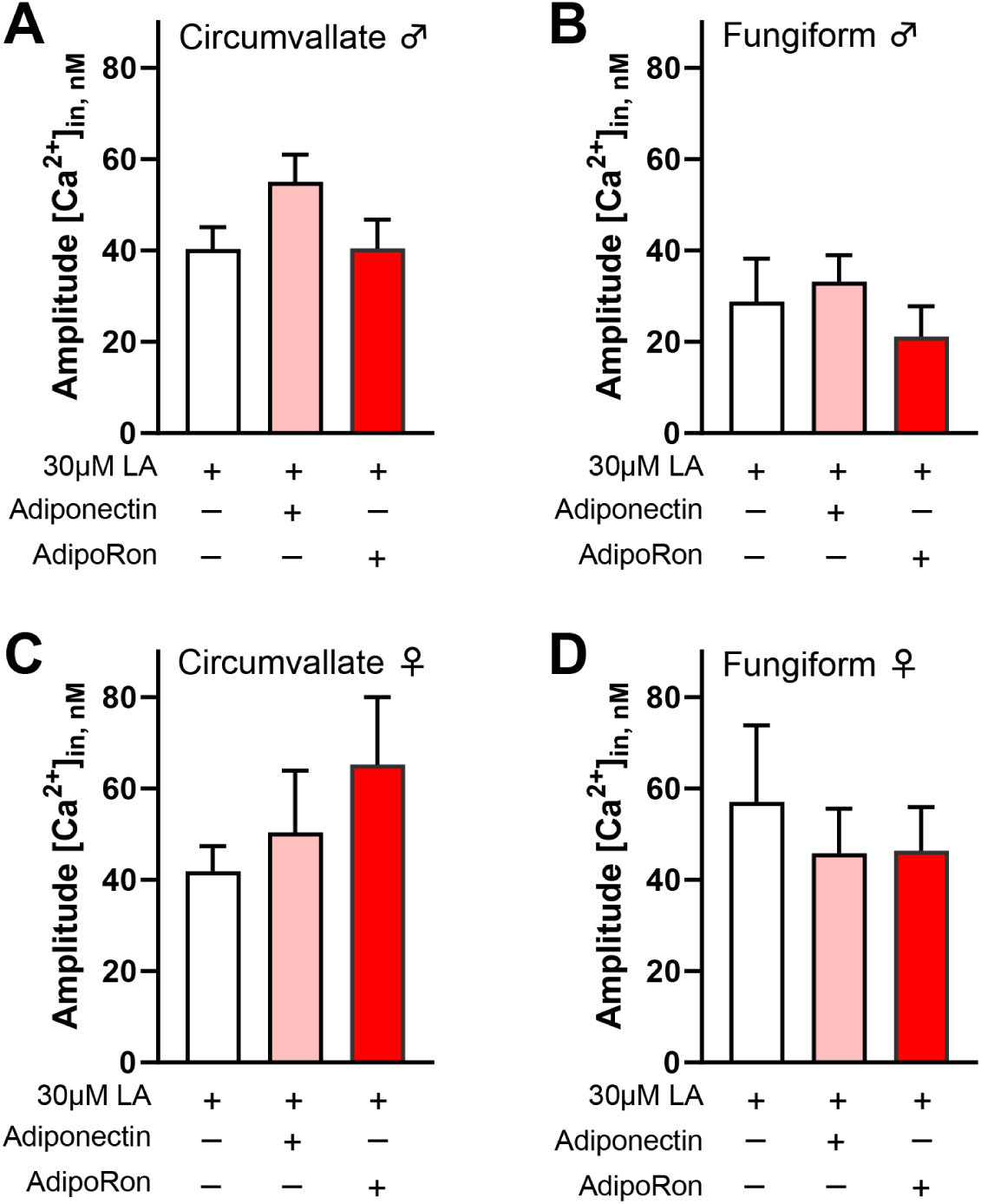
Blocking CD36 with sulfosuccinimidyl oleate (SSO) attenuates the enhancement role of AdipoRon/adiponectin on LA-induced calcium responses in mouse taste bud cells. No significant differences in calcium responses were observed in 100 µM SSO pretreated taste bud cells (isolated from (A) male circumvallate; (B) male fungiform; (C) female circumvallate; and (D) female fungiform) stimulated by 30 μM LA with or without 5 μM AdipoRon and 10 ng/ml adiponectin (P>0.05 for all tests; n = 44−71 cells from at least 3 animals for each group). Data are presented as mean ± SEM. Unpaired, two-tailed Student’s t-test (with Welch’s correction if the populations’ standard deviations are not equal) was used to determine statistical significance.

### Adiponectin/AdipoRon enhancement of LA-induced responses is dependent upon AdipoR1

Taste bud cells are known to highly express functional adiponectin receptors (Crosson et al., 2019). AdipoR1 and AdipoR2 are major receptors for the biological effects of adiponectin and are activated through AMPK and PPARα pathways, respectively. Furthermore, AMPK is required to enhance AdipoRon’s role in promoting fatty acid-induced calcium responses. Thus, the enhancement effect of AdipoRon on fatty acid-induced responses is likely to be mediated by AdipoR1, as was the case for the HuFF cells (Lin et al., 2023). To test this possibility, we performed calcium imaging to measure the cellular responses to fatty acid in isolated taste bud cells from *Adipor1*^−/−^ mice and *Adipor1*^+/+^ controls. LA-induced calcium responses in both circumvallate and fungiform taste bud cells from *Adipor1*^+/+^ mice were increased by the application of adiponectin (Figure 4A, male circumvallate, P = 0.0419; 4B, male fungiform, P = 0.0319; 4C, female circumvallate, P = 0.0499; 4D, female fungiform, P = 0.05) and AdipoRon (Figure 4A, male circumvallate, P = 0.0343; 4B, male fungiform, P = 0.0178; 4C, female circumvallate, P = 0.0042; 4D, female fungiform, P = 0.0235), but not in taste bud cells from *Adipor1*^−/−^ mice (Figure 4E-H; P > 0.05 for all tests, n = 28−100 cells from at least 3 animals for each group). Unpaired, two-tailed Student’s t-test (with Welch’s correction if the populations’ standard deviations are not equal) was used to determine statistical significance.

**Figure 4.**
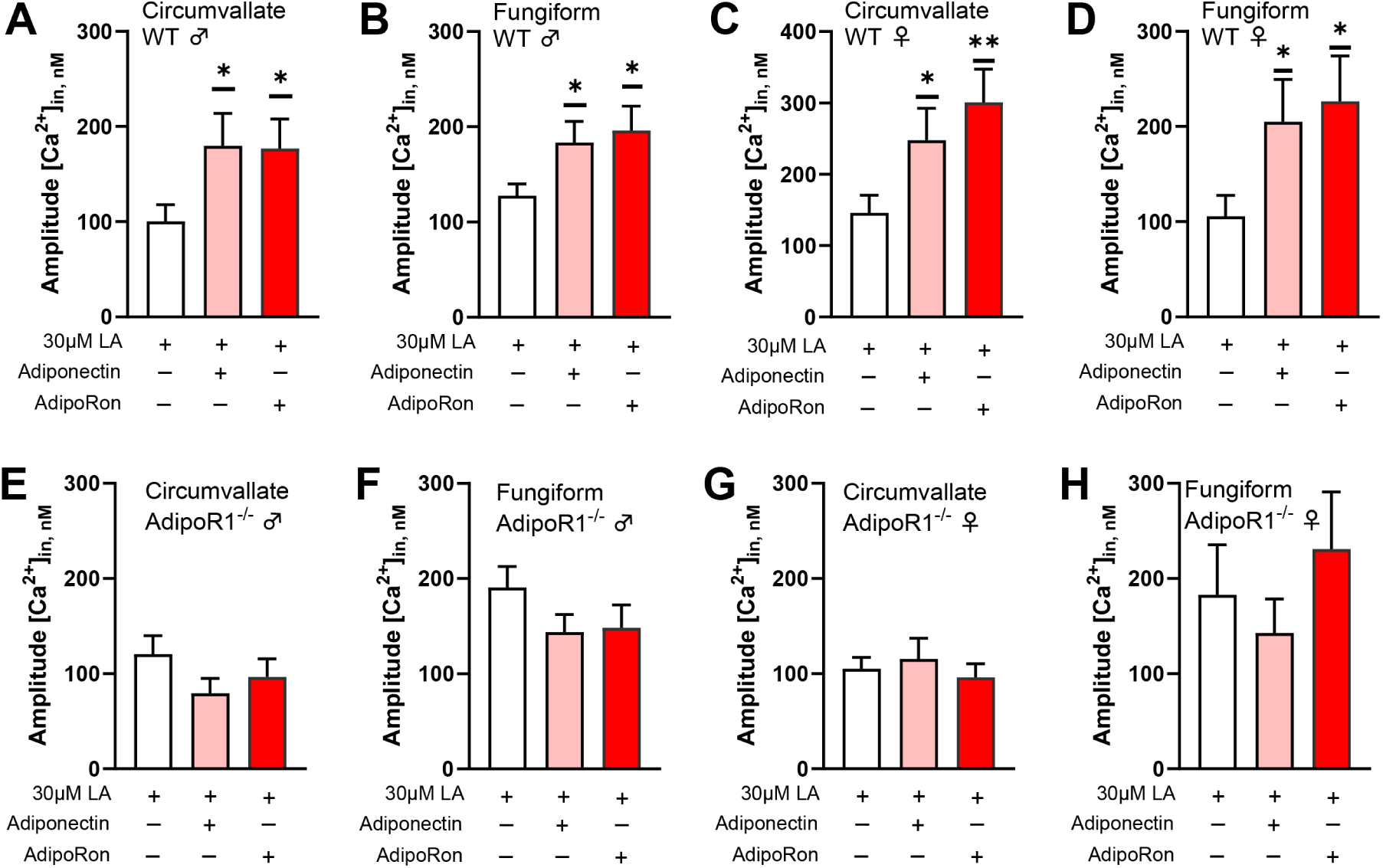
Adiponectin/AdipoRon enhancement of fatty acid responses is dependent upon AdipoR1. (A) Application of 10 ng/ml adiponectin (t_(76.57)_ = 2.069, P=0.0419) and 5 μM AdipoRon (t_(81.07)_ = 2.153, P=0.0343) was significantly increased the 30 μM LA-induced calcium responses in mouse taste bud cells from circumvallate of *Adipor1*^+/+^ males; (B) Application of adiponectin (t_(110.6)_ = 2.173, P=0.0319) and AdipoRon (t_(102.5)_ = 2.408, P=0.0178) was significantly increased LA-induced calcium responses in mouse taste bud cells from fungiform of *Adipor1*^+/+^ males; (C) Application of adiponectin (t_(89.11)_ = 1.988, P=0.0499) and AdipoRon (t_(87.25)_ = 2.938, P=0.0042) was significantly increased LA-induced calcium responses in mouse taste bud cells from circumvallate of *Adipor1*^+/+^ females; (D) Application of adiponectin (t_(77.39)_ = 1.991, P=0.05) and AdipoRon (t_(74.92)_ = 2.312, P=0.0235) was significantly increased LA-induced calcium responses in mouse taste bud cells from fungiform of AdipoR1^+/+^ females. Application of adiponectin and AdipoRon did not change LA-induced calcium responses in taste bud cells isolated from *Adipor1*^−/−^ mice (Figure 6E−H, P > 0.05 for all tests, n = 28−100 cells from at least 3 animals for each group). Data are presented as mean ± SEM. Unpaired, two-tailed Student’s t-test (with Welch’s correction if the populations’ standard deviations are not equal) was used to determine statistical significance. * p < 0.05, ** p < 0.01.

### No significant effect of AdipoRon on fat taste behavior in mice

Next, we performed brief-access taste tests to determine whether AdipoRon enhancement of fatty acid responses in mouse taste bud cells correlated with any differences in taste behavioral responses to fat. Since AdipoRon is an orally active agonist for adiponectin receptors and injection may cause stress for the animals, we tested AdipoRon’s ability to alter fat taste behavioral responsiveness in mice by adding it directly to the test solutions rather than injecting it. However, no significant genotype effect in animals’ taste behavioral responses to intralipid was found in WT and *Adipor1*^−/−^ mice with (day 3 and 4) or without (day 2 and 5) the administration of 10 μM AdipoRon (Figure 5A-D). Unexpectedly, we did observe an increase in the lick ratio for a single intralipid concentration in *Adipor1*^−/−^ males treated with AdipoRon (Figure 5C, Adjusted P=0.0333).

**Figure 5.**
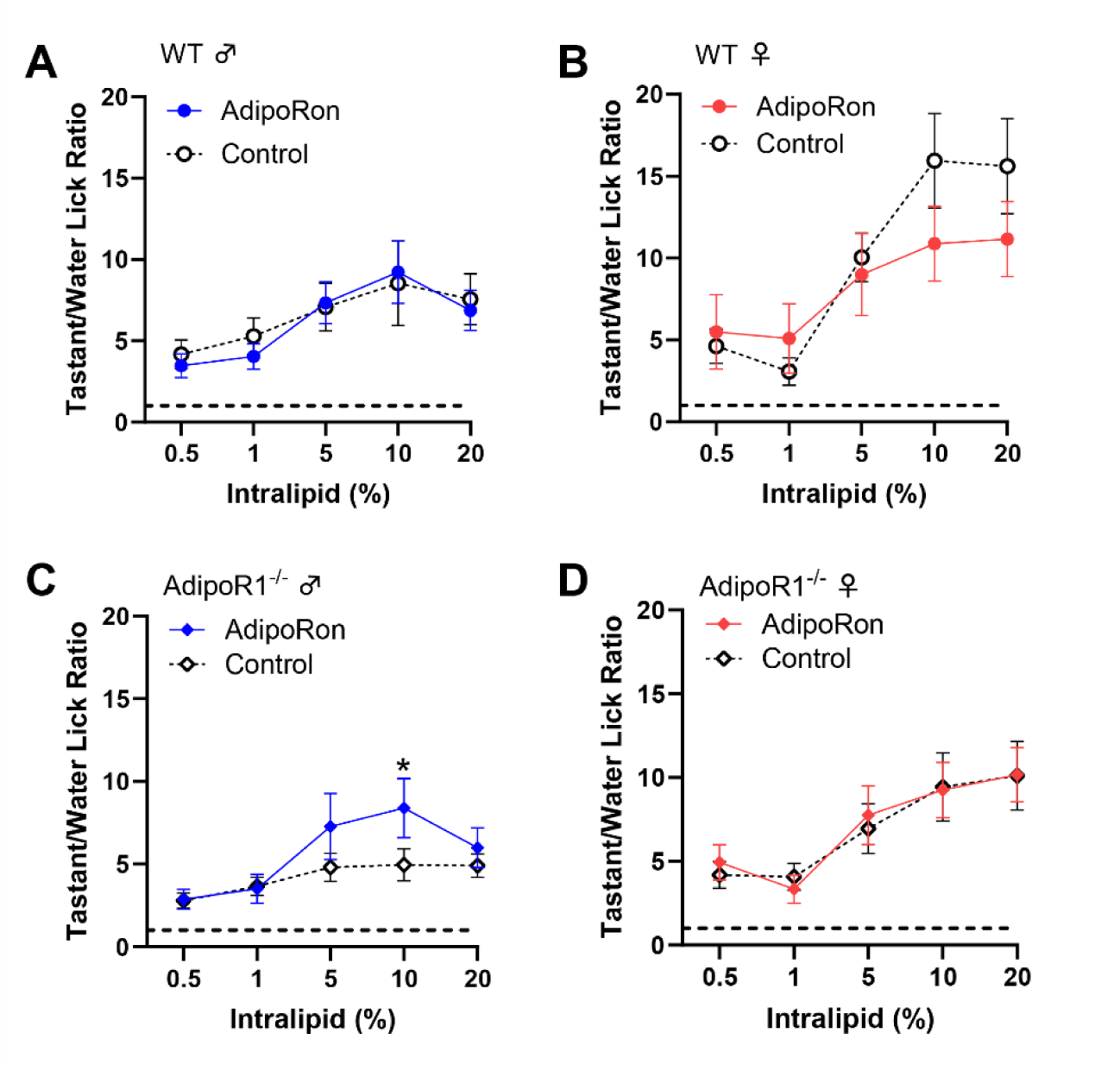
The effect of 10 μM AdipoRon on behavioral taste responses to intralipid in mice. (A) No significant difference in taste behavioral responses to intralipid was found in *Adipor1*^+/+^ males with or without the treatment of AdipoRon (F_(1, 70)_ = 0.1285, P=0.7211); (B) No significant difference in taste behavioral responses to intralipid was found in *Adipor1*^+/+^ females with or without the treatment of AdipoRon (F_(1, 70)_ = 1.256, P=0.2663); (C) An unexpected increase behavioral response to 10% intralipid (Adjusted P=0.0333) was found in *Adipor1*^−/−^ males treated with AdipoRon (F_(1, 70)_ = 3.818, P=0.0547); (D) No significant difference in taste behavioral responses to intralipid was found in *Adipor1*^−/−^ females with or without the treatment of AdipoRon (F_(1, 70)_ = 0.0212, P=0.8847). Data are presented as mean ± SEM. Two-way ordinary ANOVA with Tukey’s test for multiple comparisons was used to determine statistical significance. *p < 0.05.

### Reduction of taste behavioral responsiveness to intralipid in Adipor1^−/−^ mice

To reduce possible fat experience-induced differences in taste behavior that may affect the determination of AdipoRon’s effects on fat taste, we averaged the data from the day before (day 2) and the day after (day 5) to compare with the AdipoRon treatment days (day 3 and 4). The unexpected discovery in *Adipor1*^−/−^ males (Figure 5C) led us to compare each day’s data in the brief-access taste test more closely, especially for the results from mice lacking *Adipor1*. Interestingly, naïve *Adipor1*^−/−^ animals were indifferent to all concentrations of intralipid, and the fat taste impairment in *Adipor1^−/−^*mice does not seem to be permanent; they were able to detect intralipid after being exposed to this tastant for one or two days (Figure 6). Although both *Adipor1*^−/−^ males and females lost the ability to taste different concentrations of intralipid on the first day, they responded differently to the loss of typical fat taste. *Adipor1*^−/−^ males showed very similar licking responses to all concentrations of intralipid and pure water (Figure 7A, F_(5, 42)_ = 0.9413, P = 0.4644), whereas *Adipor1*^−/−^ females did not show any concentration-dependent licking to intralipid but could taste the difference between water and intralipid, even at its lowest concentration (Figure 7B, F_(5, 42)_ = 14.13, P < 0.0001). In contrast, the WT animals performed significantly more licks with intralipid than water in a dose-dependent manner (Figure 7C, males: F_(5, 42)_ = 4.441, P = 0.0025; 7D, females: F_(5, 42)_ = 6.036, P = 0.0003). Additionally, when *Adipor1*^−/−^ mice were initially exposed to sucrose, they could taste the difference between the tastant and water (Figure 7E-F). Our brief-access test data suggests that knocking out AdipoR1 may have a profound effect on the ability of mice to detect fatty acids.

**Figure 6.**
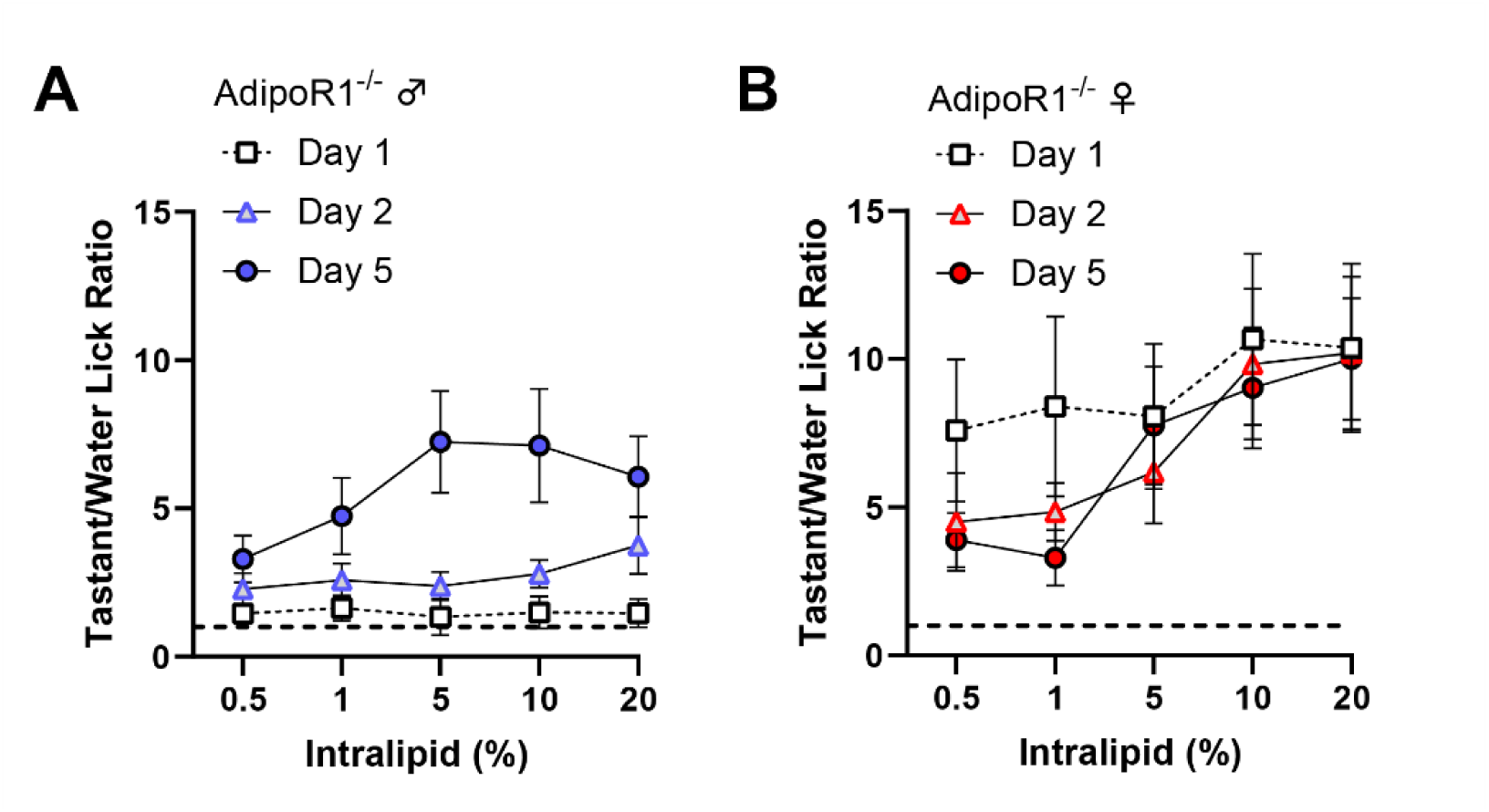
Results from brief-access taste testing of *Adipor1*^−/−^ mice differed following exposure to intralipid. (A) A significant difference in taste behavioral responses to intralipid was found in *Adipor1*^−/−^ male mice on different days (F_(2, 105)_ = 25.14, P<0.0001), *Adipor1*^−/−^ males failed to show concentration-dependent licking to intralipid on the first two days of testing; (B) No significant difference in taste behavioral responses to intralipid was found in *Adipor1*^−/−^ female mice on different days (F_(2, 105)_ = 1.524, P=0.2226). Data are presented as mean ± SEM (n = 8 for each group). Two-way ordinary ANOVA was used to determine statistical significance.

**Figure 7.**
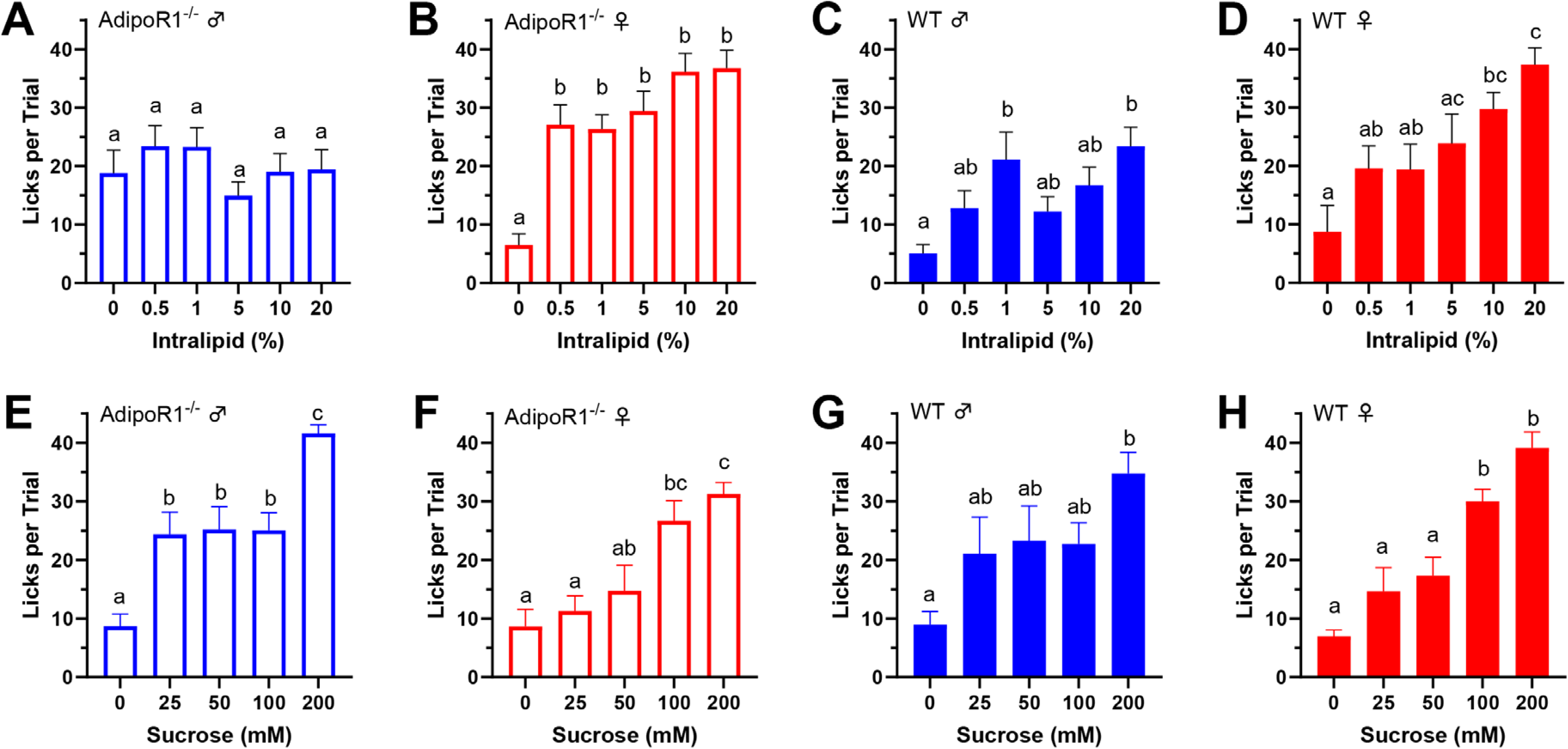
Mean number of licks per trial during brief-access tests comparing tastants and water. (A) No significant difference in linking responses to all concentrations of intralipid and pure water in *Adipor1*^−/−^ males (F_(5, 42)_ = 0.9413, P=0.4644); (B) *Adipor1*^−/−^ females did not show any difference in linking responses to all concentrations of intralipid but could taste the difference between water and intralipid (F_(5, 42)_ = 14.13, P<0.0001); (C) *Adipor1*^+/+^ males performed significantly more licks with intralipid than water in a dose-dependent way (F_(5, 42)_ = 4.441, P=0.0025); (D) *Adipor1*^+/+^ females performed significantly more licks with intralipid than water in a dose-dependent way (F_(5, 42)_ = 6.036, P=0.0003). All the animals performed significantly more licks with sucrose than water in a dose-dependent way, (E) *Adipor1*^−/−^ males (F_(4, 35)_ = 15.13, P<0.0001); (F) *Adipor1*^−/−^ females (F_(4, 35)_ = 9.901, P<0.0001); (G) *Adipor1*^+/+^ males (F_(4, 30)_ = 4.030, P=0.0099); (H) *Adipor1*^+/+^ females (F_(4, 35)_ = 20.94, P<0.0001). To better represent the real differences in taste behavioral responses between water and tastant, trials were included when the number of water licks began to drop dramatically. Data are presented as mean ± SEM (n = 8 for each group). Two-way ordinary ANOVA with Tukey’s test for multiple comparisons was used to determine statistical significance. Letters above bars indicate statistical grouping.

## Discussion

Main findings from this study support a role for adiponectin signaling in peripheral gustatory detection of fat in mice. Adiponectin/AdipoRon enhances LA-induced calcium responses in isolated WT mouse taste bud cells from both circumvallate and fungiform papillae, but this enhancement effect was not seen in taste bud cells isolated from *Adipor1*^−/−^ mice. AMPK and CD36 are required for the enhancement effect of Adiponectin/AdipoRon on LA-induced calcium responses. Although the concentration (10 μM) of AdipoRon that enhanced the cellular responses of taste bud cells to fatty acids did not seem to change fat taste behaviors in mice, we found that naïve *Adipor1*^−/−^ mice failed to taste the difference between different concentrations of intralipid.

The importance of two primary fatty acid receptors, CD36 and GPR120, in the taste perception of dietary lipids, is well established (Laugerette et al., 2005; Martin et al., 2011; Murtaza et al., 2020; Ozdener et al., 2014; Yasumatsu et al., 2019). Both CD36 and GPR120 mediate calcium responses to fatty acids in mammalian taste bud cells (Abdoul-Azize et al., 2014; Ozdener et al., 2014) and both have been implicated as critical for their contributions to fat taste (Gilbertson & Khan, 2014). Studies have shown that CD36 in other tissues can be regulated by hormones, such as GLP-2 (Hsieh et al., 2009), insulin (Luiken et al., 2002), and adiponectin (Fang et al., 2010) to stimulate lipid uptake, while GPR120 is likely to be involved in lipid-induced release of peptide hormones, such as GLP-1 (Hirasawa et al., 2005), cholecystokinin (CCK) (Tanaka et al., 2008), and pancreatic polypeptide (Zhao et al., 2020).

Given the differences in their relationships with peptide hormones, we hypothesized that the enhancement role of adiponectin/AdipoRon on fatty acid responses in taste cells is mediated by CD36. Indeed, our previous study provided evidence that the CD36 pathway, independent of the GPR120 pathway, is functionally responsible for AdipoRon’s effect on the enhancement of fatty acid-induced calcium responses (Lin et al., 2023). In detail, our results illustrated the inability of AdipoRon to enhance fatty acid-induced calcium responses in HuFF cells when pharmacologically blocking CD36 function with SSO. In contrast, AdipoRon was still able to increase the fatty acid-induced calcium responses in HuFF cells blocking GPR120 function via AH-7614, and AdipoRon was unable to affect calcium responses induced by the GPR120 agonist GW9508 (Lin et al., 2023). Studies in skeletal muscle cells and cardiomyocytes have suggested that CD36 is regulated by adiponectin signaling by two distinct ways: (1) by increasing the expression of CD36 (Fang et al., 2009); (2) by enhancing the translocation of CD36 from the intracellular to the plasma membrane (Fang et al., 2010). AdipoRon/adiponectin’s enhancement of fatty acid-induced calcium responses in taste cells occurs within several minutes. Therefore, the time course of changes in CD36 gene expression may not fit with such acute effect of AdipoRon/adiponectin in taste cells. Results from our previous research also indicated that CD36 subcellular translocation, rather than CD36 transcription, is responsible for the enhancement effect of AdipoRon on fatty acid responses in HuFF cells (Lin et al., 2023). We would expect to find similar mechanisms in mouse taste systems; however, in the present study, we focused only on the involvement of CD36 in the effect of AdipoRon/adiponectin on fatty acid-induced calcium responses in mouse taste bud cells. As expected, pharmacological blocking of CD36 function by SSO pretreatment eliminated the enhancement role of adiponectin/AdipoRon on fatty acid-induced calcium responses in taste bud cells isolated from wild-type mice, indicating the involvement of CD36 in adiponectin signaling on fat sensing in mice. Future studies are needed to address whether GPR120 signaling interacts with adiponectin signaling to influence fat taste modulation and whether adiponectin/AdipoRon stimulates CD36 translocation in mouse taste bud cells.

We also demonstrated that pharmacological inhibition of the AMPK pathway via Compound C attenuated AdipoRon’s enhancement effect on fatty acid responses in isolated wild-type mouse taste bud cells. AMPK plays an important role in regulating energy homeostasis, which mediates adiponectin-stimulated fatty acid oxidation in skeletal muscle (Yamauchi et al., 2002). Adiponectin has been reported to increase AMPK activity in the hypothalamus to stimulate food intake in mice (Kubota et al., 2007). Moreover, the presence of CD36 is required for AMPK-mediated stimulation of long-chain fatty acid uptake in cardiomyocytes (Habets et al., 2007), and AMPK promotes long-chain fatty acid uptake in intestinal epithelial cells by simultaneously regulating CD36 protein abundance and subcellular translocation (Wu et al., 2020). In addition, adiponectin-stimulated long-chain fatty acid uptake in adult cardiomyocytes was significantly attenuated by inhibition of the AMPK pathway with Compound C (Fang et al., 2010). These previous studies indicate that the AMPK-CD36 pathway is involved in adiponectin signaling. AdipoRon has been shown to increase the phosphorylation of AMPKα (Thr172) and the AdipoRon-enhanced CD36 cell surface translocation was effectively eliminated by Compound C in HuFF cells (Lin et al., 2023). Since both AMPK and CD36 are required for the enhancement role of adiponectin signaling on fatty acid responses in mouse taste bud cells, a similar mechanism may function in the mouse taste system that AdipoRon/adiponectin-stimulated activation of AMPK could lead to the CD36 translocation from the intracellular to the plasma membrane, thereby increasing fat taste responses.

Adiponectin undergoes post-translational modifications to form trimers, hexamers, high molecular weight (HMW) multimers, and super-HMW multimers (specific to saliva samples) (Lin et al., 2014; Waki et al., 2003). At least three receptors, AdipoR1, AdipoR2, and T-cadherin, have been shown to mediate the pleiotropic actions of adiponectin (Hug et al., 2004; Yamauchi et al., 2003). As one of the best-known and most abundant hormones in the plasma, the importance of adiponectin is well-established in many tissues and organs (Khoramipour et al., 2021). Adiponectin has received much attention for its role in various metabolic processes (including uptake and oxidation of lipids and carbohydrates) by stimulating the activity of AMPK and PPARα through the AdiopoR1 and AdipoR2 receptors, respectively (Yamauchi et al., 2007). Decreased adiponectin levels have been found in patients with chronic diseases, such as obesity (Arita et al., 1999), type 2 diabetes (Spranger et al., 2003), and hypertension (Adamczak et al., 2003). Interestingly, the loss of taste has also been demonstrated in such disorders (De Carli et al., 2018; Kaufman et al., 2020).

Adiponectin receptors are highly expressed in mammalian taste buds (Crosson et al., 2019), therefore it’s not hard to imagine that adiponectin signaling could play a critical role in taste regulation like other hormones, such as leptin, GLP-1, and ghrelin. However, due to the complex nature of adiponectin signaling, there is little understood about the mechanisms of adiponectin signaling modulation in the taste system. In the present study, we targeted AdipoR1 to study the role of adiponectin signaling in fat taste for several reasons as mentioned in the introduction: (1) T-cadherin is the receptor for the HMW form of adiponectin (Hug et al., 2004) and AdipoRon is an agonist of AdipoR1 and AdipoR2 (Okada-Iwabu et al., 2013), therefore T-cadherin is unlikely to be a contributor to the effects of AdipoRon on fatty acid responses; (2) the EC_50_ value of AdipoRon’s enhancement of LA-induced calcium responses (1.67 µM) (Lin et al., 2023) is close to the Kd values reported for AdipoRon binding to AdipoR1 (1.8 µM) and AdipoR2 (3.1 µM) (Okada-Iwabu et al., 2013), suggesting a potential role of these receptors in the effects of AdipoRon on fat taste responses; (3) however, AdipoR1 and T-cadherin immunolocalize to mouse circumvallate taste buds, while AdipoR2 immunolocalize to the surrounding tissues, but not in taste buds (Crosson et al., 2019); (4) adiponectin receptors mediate distinct downstream signaling pathways, AdipoR1 appears to be more tightly associated with the activation of the AMPK pathway, whereas AdipoR2 is more closely involved in the activation of the PPARα pathway (Okada-Iwabu et al., 2013; Yamauchi et al., 2007); (5) the effect of AdipoRon on LA-induced calcium responses in HuFF cells requires the activation of AMPK (Lin et al., 2023) and adiponectin failed to active AMPK in hepatocytes of Adipor1−/− mice (Yamauchi et al., 2007); (6) last but not least, a similar mechanism of adiponectin stimulates fatty acid uptake by regulating CD36 via the activation of AdipoR1 and AMPK has been demonstrated in other tissues. Combined with previously published data (Lin et al., 2023), our experiments show that adiponectin/AdipoRon enhances fatty acid-induced calcium responses in taste cells via CD36 and the increases in CD36 translocation by adiponectin/AdipoRon that occur through AdipoR1-AMPK signaling pathway (Fang et al., 2010). Therefore, we postulated a novel pathway by which adiponectin signaling modulates fat taste responsiveness in mammalian taste cells (Figure 8). Although our results indicated that AdipoR1 is the contributor to the role of AdipoRon/adiponectin on fatty acid responses in taste cells, we cannot exclude other receptors, especially T-cadherin, may also play an essential role in mammalian taste sensation. Additionally, HMW multimers have been suggested to be the major active form in mediating the multiple metabolic actions of adiponectin in periphery tissues. Due to the saliva specifically contains a super high molecular weight form of adiponectin (Lin et al., 2014) and T-cadherin is highly expressed in taste bud cells (Crosson et al., 2019), further studies are needed to fully understand the exact role of adiponectin signaling on the peripheral gustatory system.

**Figure 8.**
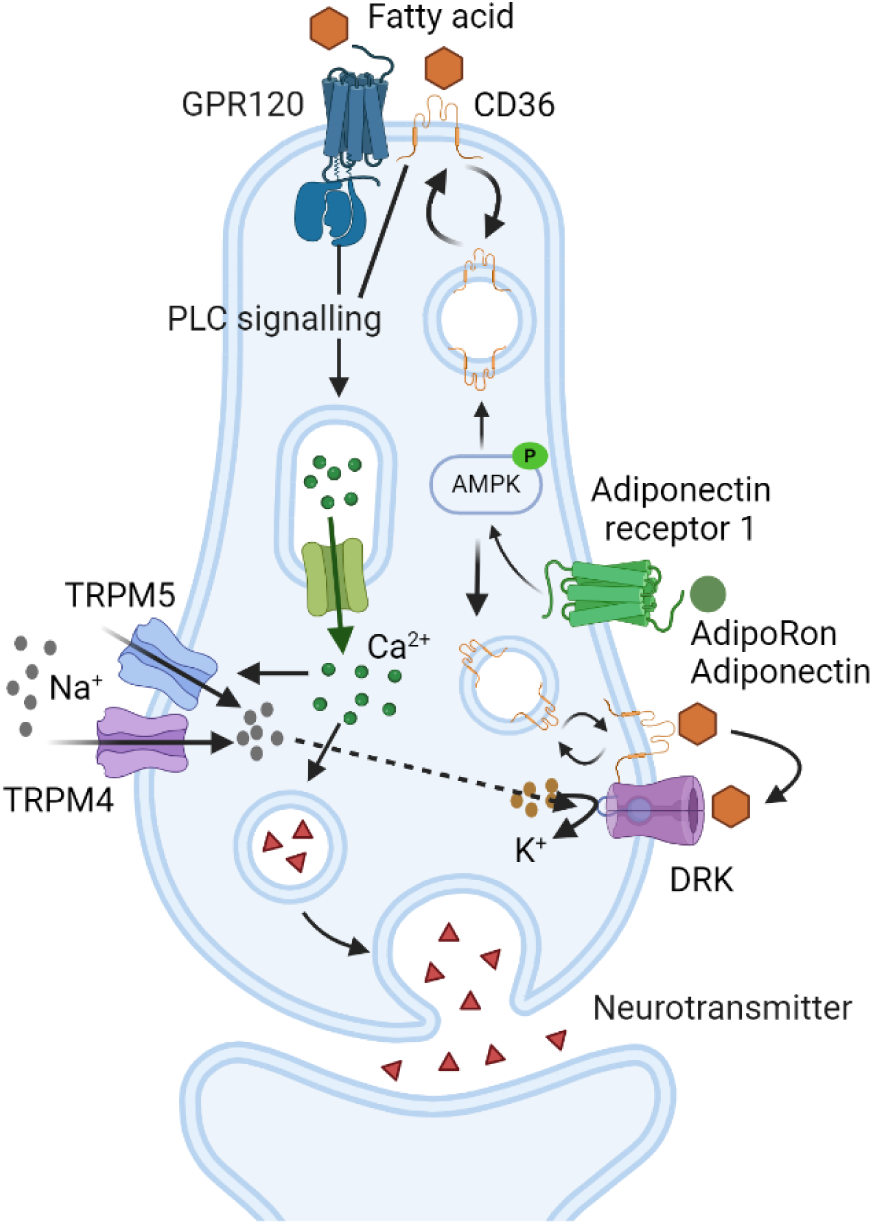
Proposed mechanism of adiponectin signaling on fat sensing in taste cells. Adiponectin (or AdipoRon) binds to adiponectin receptor 1 (AdipoR1), which stimulates the activation of 5’ adenosine monophosphate-activated protein kinase (AMPK). In response to the stimulation with adiponectin (or AdipoRon), cluster of differentiation 36 (CD36) translocates from intracellular storage depots to the cell membrane to increase fatty acid taste signaling. Fatty acids bind to their G protein-coupled receptors on the cell membrane, release G proteins, stimulate phospholipase C-beta 2 (PLCβ2), generate second messengers, elicit an increase in intracellular calcium, activate transient receptor potential channel M4 and/or M5 (TRPM4/TRPM5), induce cell depolarization. CD36 and fatty acids may interact directly with PLCβ signaling, or CD36 may facilitate fatty acids binding with the receptors (GPR120) to enhance the taste cell activation or block fatty acid-sensitive delayed rectifying K^+^ (DRK) channels to prolong depolarization of the cell.

Our results showed that adiponectin/AdipoRon enhances fatty acid-induced calcium responses through AdipoR1-AMPK-CD36 signaling in taste cells, and we hypothesize that AdipoRon plays a physiological role in the modulation of fat taste. We used brief-access taste testing assay to test the effects of AdipoRon on behavioral responses to intralipid in WT and *Adipor1*^−/−^ mice. However, in our experiments, AdipoRon concentration (10 μM) that enhances the cellular responses of taste bud cells to fatty acids does not appear to alter the intralipid/water ratio in brief-access testing in mice. It is worth noting that the cell-based experiments were performed *in vitro* in the absence of endogenous adiponectin, whereas the behavioral studies were conducted in animals rich in salivary and circulating adiponectin. It has been reported that adiponectin knockout mice and WT mice have equivalent taste behavior responses to different taste stimuli, including intralipid; and the salivary gland-specific adiponectin rescue, but not the global adiponectin rescue in adiponectin KO mice significantly increases behavioral taste responses to intralipid (Crosson et al., 2019). Therefore, our results are not particularly surprising because AdipoRon’s effect is likely masked by endogenous adiponectin, especially salivary adiponectin. Thus, adiponectin KO mice, without affecting endogenous adiponectin, should be an effective animal model to verify the *in vivo* effect of AdipoRon on fat taste. Moreover, it is also possible that the concentrations we used were insufficient to elicit differences in taste behavior or that the drug delivery method we applied was inappropriate to induce behavioral changes in taste. Higher concentrations of AdipoRon or other drug delivery approaches are needed to elucidate the functional role of AdipoRon in regulating fat taste.

No significant differences were found in the taste behavioral assay in WT animals treated with AdipoRon. If anything, there was a trend toward AdipoRon in female mice to alter the licking responsiveness in brief-access taste test. A recent study reported that male but not female diabetic mice were resistant to globular adiponectin (gAd) treatment, and that gAd-treated female but not male mice displayed higher AMPK activity and CD36 protein levels compared to controls (Chattopadhyay et al., 2022). The adiponectin levels are higher in females than males (Böttner et al., 2004; Song et al., 2014). Recently, it was shown that differing adiponectin concentrations may account for the sex difference in human olfactory sensitivity (Pfabigan et al., 2022). Data from our lab showed that females had higher fat taste sensitivity and could detect lower concentrations of fatty acids (Dahir et al., 2021). It is still unclear if and how adiponectin signaling may influence fat taste detection ability in sex-dependent way. Further studies are needed to explore whether the sex differences in fat taste sensitivity and in adiponectin concentrations are related.

Although our experiments could not provide direct evidence that AdipoRon affects fat taste behavior, the data from brief-access testing showed that fat naïve *Adipor1*^−/−^ animals were indifferent to all concentrations of intralipid, suggesting adiponectin signaling has a profound effect on the ability of mice to detect fatty acids. Unlike *Adipor1*^−/−^ females performed more licks per trail with intralipid than pure water, *Adipor1*^−/−^ males did not show any significant difference in licks comparing intralipid and water, indicating a sex-dependent role of adiponectin signaling on fat taste behavior or a sex-dependent response strategy in the face of fat taste loss. Knocking out of *Adipor1* may alter critical downstream cellular signaling in different ways between males and females, and there may be compensatory mechanisms that is able to supplement for the lack of adiponectin signaling in females. Our cell-based experiments showed that adiponectin signaling enhances the cellular responses to fatty acids in taste cells. We expected that *Adipor1* deficiency would have reduced behavioral responses to fat stimuli in mice, however the completely abolished taste behavioral responses to intralipid in *Adipor1*^−/−^ animals in our experiments was unexpected, since these mice should contain intact fat taste signaling components. The reduction of CD36 on the tongue has been shown to attenuate linoleic acid preference in obesity-prone and obesity-resistant rat (Chen et al., 2013). Similarly, CD36 knockout mice failed to prefer to fat emulsion in both short and long term two-bottle tests (Laugerette et al., 2005) and the fat preference was fully abolished not only in CD36 knockout mice but also in the heterozygous CD36^+/−^ animals (2-fold lower CD36 protein level in circumvallate papillae) (Martin et al., 2011). Recently, it was reported that CD36 protein levels were significantly reduced (3.7-fold) in the retina of *Adipor1-*deficient mice (Lewandowski et al., 2024). Our behavioral studies demonstrated that fat taste loss was found only in *Adipir1* knockout mice but not in WT animals and the *Adipir1* knockout animals were indifferent to intralipid but not for sucrose. AdipoRon/adiponectin activates AMPK through AdipoR1, increases the translocation of CD36 from intracellular to the cell membrane, thereby selectively enhancing the cellular responses to fatty acids. Therefore, lower CD36 protein levels in taste bud cells, if true, combined with the lack of adiponectin signaling to stimulate CD36 translocation to maintain certain cell surface expression levels, may contribute to the absence of fat behavioral responses in *Adipor1-*deficient mice.

After being exposed to the intralipid for one (females) or two (males) days in the brief-access test, *Adipor1*^−/−^ mice were able to detect the different concentrations of intralipid. CD36 gene expression and protein levels in mouse circumvallate papillae were sensitive to the dietary lipid content and affected by the diet intake during a fasting/re-feeding sequence (Martin et al., 2011). Therefore, changes in CD36 levels induced by lipid exposure may contribute to the recovery of taste ability to fatty acids. Besides, we cannot exclude the possibility of the post-oral nutritive effects attribute to the taste recovery. Experience-induced recovery of fat preference has been found in taste-deficient animals with genetic knockout of gustatory fat taste signaling components, such as CD36, CALHM1, TRPM5, and P2X2/P2X3, and these animals could learn to use residual oral fat sensory cues based on post-oral reinforcing actions to display strong preferences and high intakes of intralipid (Sclafani & Ackroff, 2014, 2018, 2022; Sclafani et al., 2007). Further studies are needed to better understand the role of adiponectin signaling in fat taste and to explore the links between adiponectin signaling, fat taste, dietary fat intake, and obesity.

## Author Contributions

F.L. and T.A.G. conceived and designed research; F.L. and E. M. performed experiments; F.L. analyzed data; F.L. and T.A.G. interpreted results of experiments; F.L. and T.A.G. prepared figures; F.L. prepared first draft of manuscript; F.L., E.M., and T.A.G. edited and revised manuscript; F.L., E.M., and T.A.G. approved latest version of the manuscript; T.A.G. secured funding.

## Institutional Review Board Statement

All procedures involving animals were approved by the Institutional Animal Care and Use Committee of the University of Central Florida (protocol 2023-84; approved most recently on 4 June 2023) and were performed in accordance with American Veterinary Medical Association guidelines.

## Data Availability Statement

All relevant data are within the manuscript.

