## Supplemental Figure S1 for "Adiponectin signaling modulates fat taste responsiveness in mice" (from another version of this preprint: version 1, 2024-09-06)

**Supplementary Material**

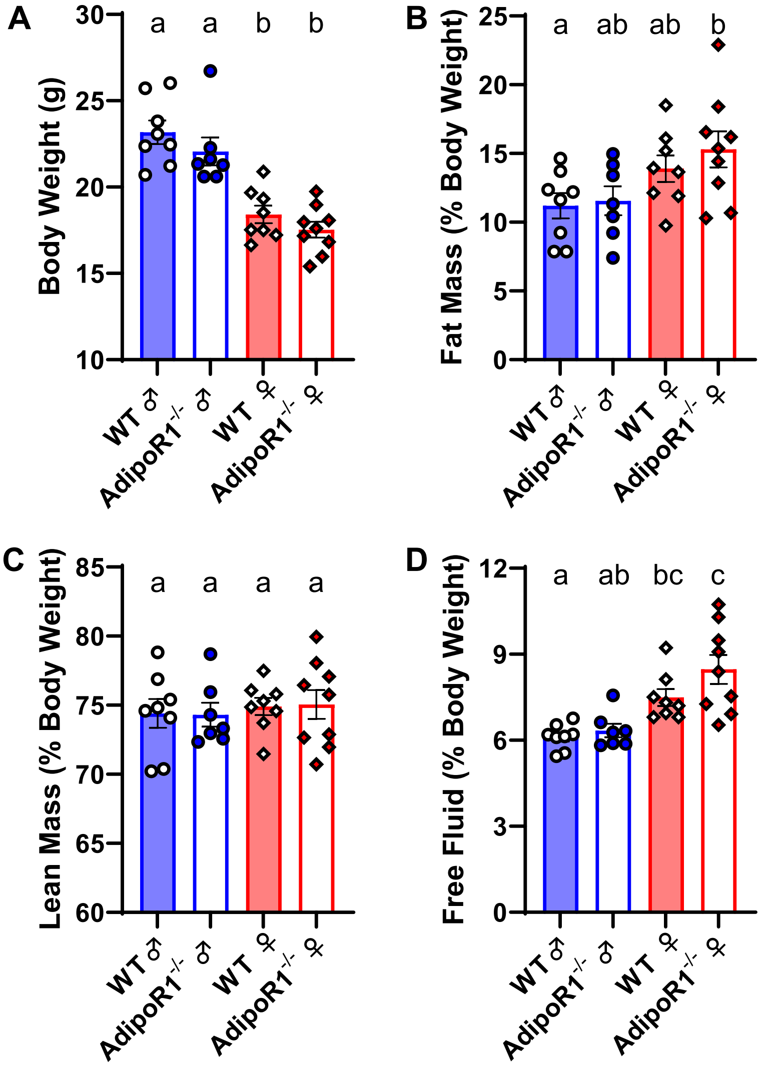


Figure S1. Body weight and total body composition in AdipoR1$^{+/+}$ and AdipoR1$^{-/-}$ mice. (A) There was no significant genotype effect on the body weight of animals at about six weeks of age (F (1, 28) = 2.655, P=0.1144), but there was a sex difference (F (1, 28) = 57.76, P<0.0001); (B) There was no significant genotype effect on the percentage of fat mass in animals (F (1, 28) = 0.646, P=0.4281), but there was a sex difference (F (1, 28) = 8.633, P=0.0065); (C) No significant difference was found in the percentage of lean mass among the animals (genotype effect: F (1, 28) = 0.0008), P=0.9774, and gender effect: F (1, 28) = 0.4542, P=0.5059); (D) There was no significant genotype effect on the percentage of free fluid in animals (F (1, 28) = 3.014, P=0.0936), but there was a sex difference (F (1, 28) = 25.53 , P<0.0001). Data are presented as mean ± SEM (n = 7−9 mice for each group). Two-way ordinary ANOVA with Tukey's test for multiple comparisons was used to determine statistical significance. Letters above bars indicate statistical grouping.
